# Modulation of slow and fast oscillations by direct current stimulation in the cerebral cortex in vitro

**DOI:** 10.1101/246819

**Authors:** Mattia D’Andola, Julia F. Weinert, Maurizio Mattia, Maria V. Sanchez-Vives

**Affiliations:** Systems Neuroscience, Institut d’Investigacions Biomèdiques August Pi i Sunyer (IDIBAPS), Barcelona, Spain; IstitutoSuperiore di Sanità, Rome, Italy; Institució Catalana de Recerca i Estudis Avançats (ICREA), Barcelona, Spain

**Keywords:** electric fields, cerebral cortex, slow oscillations, Up states, Up and Down states, beta oscillations, gamma oscillations, kainate, electrical stimulation, cortical stimulation, excitability, emergent activity, emergent properties, direct current stimulation

## Abstract

Non-invasive brain stimulation techniques, such as transcranial direct current stimulation (tDCS), play a growing role in the treatment of neurological disorders. However, the mechanisms by which electric fields modulate cortical network activity are only partially understood. To explore the spatiotemporal modulation of cortical activity by electric fields (DC fields), we exposed neocortical slices to constant fields of varying intensity and direction and we measured their effect on the low (<1 Hz) and high frequencies (beta 15-30 Hz and gamma 30-90 Hz) of spontaneously generated cortical oscillations. Slow oscillations consist of Up (active) and Down (silent) states. We found that DC fields ranging from -6 to +6 V/m induced an exponential increase in the frequency of slow oscillations through the regulation of the excitability and duration of Down states, while hardly affecting Up states duration. A computational model based on the mean-field theory of attractor dynamics provided a mechanistic and quantitative description of the network dynamics underlying such precise modulation of slow oscillatory frequency. The modulation of high frequencies by DC fields was less consistent, the high frequency power varying with the intensity of the fields only in a fraction of slices. Interestingly, negative DC fields of increasing intensities progressively and effectively reversed the increase in high frequency power induced by kainate application. Our findings have implications for the understanding of cortical oscillations and the mechanisms by which they are modulated by DC fields and may contribute to the future development of tools with an accurate spatiotemporal control of cortical activity.

**Significance statement:** Acting on the brain through electrical stimulation in order to correct dysfunctions or to induce functional recovery is a relatively common technique nowadays used in the clinical realm. In spite of the existence of previous studies on the effect of electric fields on neuronal and network physiology, questions regarding the mechanisms underlying exogenous electrical modulation of cortical dynamics still remain open. We demonstrate that continuous electric fields between -6 and +6 V/m induce a precise modulation of slow and fast cortical rhythms. Based on both experimental evidence and theoretical analysis, we describe some of the mechanistic underpinnings at play and provide useful information for the development of tools with better spatiotemporal control of cortical activity.

## Introduction

Transcranial direct current stimulation (tDCS) is a brain stimulation technique that involves the application of constant low-intensity currents between two electrodes placed on the skull. It has been reported that this type of stimulation can be effective for treatment in a wide range of neuropsychiatric disorders [1–3], in stroke rehabilitation (for a review, [4]) as well as for the improvement of memory and motor performance in healthy subjects [5, 6]. A broader range of applications and regulatory considerations were recently reviewed by an expert panel [7]. However, these findings remain somewhat controversial [8, 9] and, despite the relatively widespread use of this technique in clinical settings, the detailed mechanisms underlying the effects of DC stimulation on the cortical circuits remain to some extent unknown.

Nonetheless, several studies in animals have provided insights into the underlying mechanisms of tDCS, extensively reviewed in [10]. *In vitro* studies on rat and guinea pig hippocampal slices have shown that the application of DC fields can either depress or potentiate population spiking activity by respectively hyperpolarizing or depolarizing the neuron membranes depending on the direction of current flow [11, 12]. Similar effects were observed in rat hippocampal pyramidal neurons, where the authors showed that the degree of polarization highly depends on the relative orientation of the electric fieldswith respect to the neurons and their morphology [13], facilitating LTP [14] and modifying the global network dynamics [15]. On the other hand, fewer studies have focused on the effect on the cerebral cortex *in vitro*, although tDCS is used and mostly effective on the surface of the brain, i.e. the cortex. The application of DC fields for long periods (60 s), with intensities similar to the endogenous field potentials (0.5 - 4 V/m, [16, 17]), enhanced cortical activity in acute ferret cortical slices [18]. In that study, the weak but global polarizationof the neuron membrane voltage introduced by AC, pulsed electric fields were sufficient to entrain the macroscopic network dynamics, more specifically the cortical slow oscillations [18].

Slow oscillations typically dominate the neocortex during non-rapid eye movement (non-REM) sleep and deep anesthesia and occur at a frequency of < 1 Hz, being characterized by the alternation of active Up states and silent Down states [19]. During Up states, the network sustains persistent neural activity with a characteristic temporal structure that shares features with the awake states, such as synchronization in beta (15-30 Hz) and gamma (30-90 Hz) frequencies [20]. Slow oscillations are an emergent activity from the cortical network, therefore integrating intrinsic, synaptic and connectivity properties of both excitatory and inhibitory neurons which determine the features of the resulting spontaneous pattern of activity. Slow oscillations have been proposed to be a default activity of the cortical network [21, 22] since it is generated even when the region in which it is detected is functionally or physically disconnected from the rest of the brain, implying that this activity mostly relies on local connectivity [23].

Another motivation to study slow oscillations and the effects of electric stimulation originates from reports showing the regulatory effects of electric fields on slow waves during sleep (e.g. [6] but see[24]). It has been reported that electric stimulation can induce slow waves. In humans, tDCS and transcranial magnetic stimulation (TMS) increased slow wave activity during sleep and this resulted in the potentiation of memory performance [6, 25–27] and synaptic plasticity [28]. Therefore, because of the potential of studying them in cortical slices, their modulation by electrical stimulation, the view that they may represent default cortical activity and the promising benefits of modulating them in humans, slow oscillations provide a paradigmatic model to explore the impact of electric fields on cortical activity.

Based on the available evidence and the missing gaps regarding the mechanisms underlying the effects of electric stimulation on cortical activity, we then studied the effects of DC fields of varying intensities and directions on the emergent cortical activity, focusing both on the slow oscillations and on the beta/gamma frequency bands that occur during the Up states of these slow oscillations. In parallel, we devised a cell-assembly model based on the mean field-theory of attractor dynamics [29] to further investigate the multi-scale phenomena observed in the experiments. We observed a strong modulation of spontaneously occurring, physiological cortical oscillations with DC fields and explored both experimentally and theoretically their mechanistic underpinnings. These findings bring insight into the network effects of DC fields and their therapeutic applications.

## Material and methods

### Experimental procedures

#### Slice preparation

All animal experiments were approved by the Ethics Committee of Hospital Clinic (Barcelona, Spain). Ferrets were cared for and treated in accordance with Spanish regulatory laws (BOE-A-2013-6271), which comply with the European Union guidelines on protection of vertebrates used for experimentation (Directive 2010/63/EU of the European Parliament and the Council of 22 September 2010).

Slices were prepared as previously described [30]. Briefly: ferrets (2-7 months old, either sex) were anesthetized with sodium pentobarbital (40 mg/kg) and decapitated. The entire forebrain was rapidly removed and transferred to oxygenated cold (4-10 °C) bathing medium. Coronal slices (400 μm thick) of the occipital cortex containing primary and secondary visual cortical areas (areas 17, 18, and 19) or of the prefrontal cortex were used. A modification of the sucrose-substitution technique developed by [31] was used during the preparation to increase tissue viability.

Slices were then placed in an interface style recording chamber (Fine Science, Foster City, CA), and bathed for 15 min in an equal mixture of the sucrose-substituted solution and Artificial Cerebro Spinal Fluid (ACSF). Throughout the rest of the experiment, an in vivo-like modified ACSF was used [23]. ACSF contained (in mM): NaCl, 126; KCl, 2.5; MgSO_4_, 2; NaH_2_PO_4_, 1; CaCl_2_, 2; NaHCO_3_, 26; dextrose, 10, and was aerated with 95% O_2_, 5% CO_2_ to a final pH of 7.4. The in vivo-like modified ACSF had the same ionic composition except for different levels of (in mM): KCl, 4; MgSO_4_, 1 and CaCl_2_, 1. The temperature during the experiment was maintained at 34.5-36 °C. Electrophysiological recordings started after allowing at least 2 h of recovery.

#### Electrophysiological recordings

Extracellular local field potential (LFP) recordings from deep cortical layers (mostly layer 5) were obtained with 2-4 MΩ tungsten electrodes (FHC, Bowdoinham, ME). The raw signal was amplified by 1000 (NeuroLog System, Digitimer, Letchworth, UK) and digitized at 5 kHz (Power1401 ADC/DAC, Cambridge Electronic Design, Cambridge, UK).

#### DC electric fields and experimental protocols

A uniform DC electric field was generated as previously described [12, 18]. In short: current was passed between two customized silver/silver-chloride wires (1 mm diameter, 10 mm length) aligned in parallel (Fig. 1A). They were placed, depending on the slice size, 5-8 mm apart such that the electric field was oriented perpendicular to the wires. The cortical slices were placed such that the electric field would be parallel to the apical-dendritic axis of cortical pyramidal cells (Fig. 1A). The stimulation was timed using a Power1401 ADC/DAC (Cambridge Electronic Design, Cambridge, UK) and converted to current using a stimulus isolator (360A, WPI, Aston, UK). The applied electric fields were measured and calibrated before every experiment. In a separated test, we performed a systematic measure of the voltage gradient along the overall recording area to assure homogeneity of the electric field generated by our electrodes. Increasing fields (+/-1 to 6 V/m) were applied for 150 s, with 150 s of recovery between stimuli. Positive fields were defined as fields oriented from the white matter to the cortical surface, and negative fields were oriented in the opposite direction, thus inducing depolarization or hyperpolarization of pyramidal neurons, respectively [13, 32]. Positive and negative fields ranged from -6 to +6 V/m and were alternated to avoid tissue and electrode damage, thus progressing from +1 to -1 to +2 to -2 V/m and so on, until ±6 V/m.

**Figure 1.**
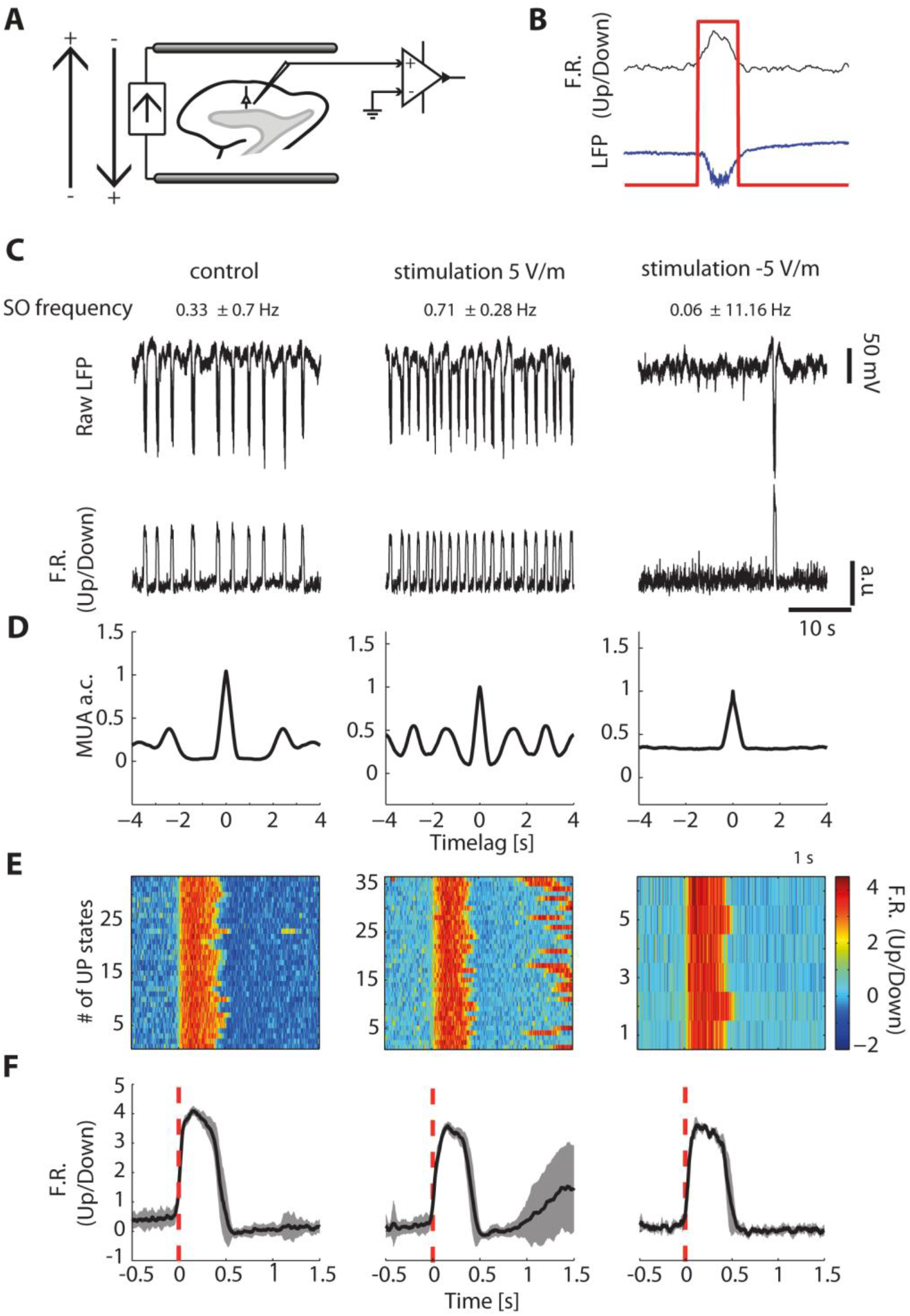
Experimental setup and response to stimulation in a representative slice. **A.** Scheme of electrode setup indicating orientation of DC fields and recording area. **B.** Example of one representative Up state showing the recorded local field potential (LFP; blue trace), the relative firing rate (F.R.; black trace) and the detection of the Up state by the used algorithm (red line) [42]. **C-F.** Representative recording from 1 slice in control conditions (left), after +5 V/m (center) and after -5 V/m DC fields (right). The average slow oscillation (SO) frequency for each condition is reported on top. **C.** Raw LFP of the signal (top) and relative firing rate from which Up states were detected (bottom). **D.** Autocorrelation (a.c.) of the multiunit activity (MUA) waveforms, considering a maximum time lag of 4 s. **E.** Raster plot of the relative firing rate (i.e. Up/Down firing rate; color coded) corresponding to all Up states evoked during 100 s of recording aligned to their onset (t=0 s). Notice that the number (#) of Up states varied with the DC field intensity. The time scale is the same as in F. **F.** Mean relative firing rate for all Up states displayed in the raster plots in E.

Two different experimental protocols were used. First, for n=21 slices, electric fields were applied during spontaneous oscillatory activity. Then, for a second group of n=15 slices, bath application of 200 nM kainate (Tocris, Bristol, UK) was used to enhance activity in the beta and gamma frequency range before applying electric fields. Pharmacological and electrical stimulation were applied only after a 300-s control recording of spontaneous activity was obtained.

### Computer model

To further investigate the mechanisms underlying the response of a cortical network generating slow oscillations to the application of constant electric fields, we devised a cell assembly computer model composed of 4000 excitatory (80%) and 1000 inhibitory (20%) leaky integrate-and-fire neurons, based on the mean field theory of attractor dynamics. The model’s theory is explained in detail in [29]. The computational network displayed slow oscillations that alternated between high-(Up) and low-firing rate (Down) states (Fig. 5A-C). Briefly, the average firing rate ν of a neuron can be modeled as a function of the instantaneous mean and variance of ionic currents flowing across the neuron’s membrane [33]. Provided that the statistics of such currents is similar for different neurons, the same input-output relationship can be used to describe the behavior of a neuronal population *(mean-field approximation)* [33–35], allowing to predict the population firing rate of a network of spiking neurons under stationary conditions throughout the computation of an input-output gain function Φ(ν) [33] that we derived as in previous models [36]. The difference between the output and the input firing rate of the network determines an effective force driving the network activity. A positive difference brings the neurons to fire at a higher rate, while a negative difference lowers their firing rate. When the input and output activity are equal, the difference is 0 and this results in null forces and the network getting stuck in a fixed point. If forces nearby such state are “attractive”,-they tend to bring the network to these states-, these preferred activity levels are called “attractor states” [29]. An activity-dependent modulation of the attractor dynamics was inserted in our model by using a self-inhibition variable proportional to a fatigue level c(t) (see below - Fig. 5A) to mimic the real time spent in each attractor (Fig. 5C) and reproduce the regularity of the slow oscillations observed in our experiments [29, 37–39]. The interplay between the non-linear dynamics of *ν*(t) and the fatigue, brings the network to a “relaxation oscillator” behavior, fitting our experimental evidence.

From the instantaneous frequency of spikes (firing rate) emitted by the whole network, an *in silico* multiunit activity (MUA) was worked out by adding a relatively weak white noise to emulate unspecific background fluctuations occurring in experimental recordings (Fig. 5A). Up states occurred in 25% of the modeled excitatory cells, resulting in sparse oscillatory activity similar to what has been reported in experiments [40, 41]. In the following paragraph the model parameters and the simulation of DC fields will be explained in detail.

### Data analysis

In order to study modulatory effects of DC fields on temporal parameters of cortical slow oscillations we compared standard slow oscillation parameters (i.e. Up and Down state duration, slow oscillation frequency, firing rate, high frequency content) in control conditions and when applying DC electric fields for varying intensities and directions. We then compared the results between each DC condition (different intensities) to the control conditions for each parameter. All analysis was performed with custom software written in Matlab (Mathworks, Natick, MA).

#### MUA computation

To detect the slow oscillation and its components, LFP recordings were first notch-filtered and de-trended to eliminate ultraslow oscillations (<0.5 Hz) by subtracting the moving average (window length 2 s). Multiunit activity (MUA) was then estimated to obtain the population firing rate as previously described [42]. In brief, given that power spectra of population firing rate have Fourier components proportional to the firing rate itself [43], high frequency components of the LFP can be seen as a linear transform of spiking activity (Fig. 1B). Thus, power changes in the Fourier components at high frequencies of the LFP provide a reliable estimation of the population firing rate. MUA traces were calculated as the average power of the normalized spectra in the frequency band from 200 to 1500 Hz. To balance large fluctuations of nearby spikes, MUAs were scaled logarithmically. Furthermore, a moving average with a sliding window of 80 ms was applied to smooth the log(MUA) time series. Since this is a relative measure resulting from an average of power spectral ratios, the log(MUA) is referred to as “relative firing rate” [42] or “network firing rate” (Fig. 1B).

#### Up and Down state detection

Up and Down states were next detected by setting time and amplitude thresholds in the log(MUA) (see Fig. 1B for an example). Up and Down state durations were computed respectively as the time between the onset and the offset of an Up state and the time between the offset of an Up state and the onset of the following one. The frequency of the slow oscillation (SO frequency) represented the duration of a complete Up-Down cycle, i.e. the time between the onset of an Up state and the onset of the following one.

#### The Modulation Index

We calculated a Modulation Index as the slope of the linear regression between the logarithm of the Down state durations and the different DC electric fields intensities (see Fig. 3A). The Modulation Index was then used to identify and exclude slices which were not modulated. We chose the lower 95 percentile as limit, i.e. slices that showed a Modulation Index lower than 0.05 were excluded from these averages as their Down states were not modulated.

#### Modulation of temporal parameters

A set of measurements was obtained to study changes in the firing rate of the cortical network. First, we analyzed changes in the relative firing rate for the different electric field intensities. By aligning all Up states to their onset (Fig. 1E) we were able to compute the mean relative firing rate for each condition as the mean value between the onset and the offset of the averaged log(MUA) waveform (Fig. 1F) of all Up states. In addition to the relative firing rate (i.e. the firing rate of the Up state versus the firing rate of the Down state), we were also interested in a modulation of the absolute firing rate during the Up state and the Down state. To do so we computed the averaged power in the range of 200 – 1500 Hz for each electric field intensity first for only Up states and then for only Down states. For the same purpose of identifying changes in the network firing, we also computed the standard deviation of the relative firing rate for Down states, averaging the log(MUA) of each Down state value between 250 ms and 50 ms before the following Up state initiated. All parameters were averaged across all Up and Down states occurring during the period of a specific condition and additionally, for each slice, all data was normalized to the control condition (i.e. without stimulation or before drug application) of this respective slice.

#### Analysis of high frequencies

Power spectrum density (PSD) analysis was carried out by Welch’s method with Hanning windowing to analyze changes in high frequency (beta and gamma) activity taking place during Up states. Changes in beta (15-30 Hz), low-gamma (30-60 Hz) and high-gamma (60-90 Hz) frequency ranges were studied by filtering the raw LFP signal using a Butterworth band-pass filter (order 3; Matlab butter function). For the same purpose, spectrograms were computed from the downsampled (1 kHz) LFP signals (Matlab spectrogram function, Hanning windowing). The spectrograms were then smoothed with a rotationally symmetric Gaussian low-pass 2D filter (size = 5, sigma = 3; matlabfspecial function) to improve clarity of the plots. Averaged power of the beta, low-gamma and high-gamma ranges (the last two sometimes unified in the total gamma) was computed from the single PSDs, normalized to the control condition and averaged across different slices, to compare the frequency behavior in each of the different conditions.

#### Statistics

A one-sample Student t-test was used to evaluate changes in average Up and Down state duration, SO frequency, relative and absolute firing rate and averaged power spectrum in the beta and gamma ranges during DC stimulation and after the application of kainate. Differences on average power induced by stimulation with respect to the averaged power of the kainate condition were studied with a two-sample Student t-test.

All mean values are represented together with their respective standard errors (SE). Statistical tests applied to data in figures have been included in the figure legends.

### Spiking neuron network model and theory

*In vitro* neuronal activity was modeled by resorting to a network of 4000 excitatory (E) and 1000 inhibitory (I) LIF neurons, having membrane potential *V(t)* dynamics d*V*/d*t* = –*V*/τ +*I/C*, with decay constant τ of 20 ms and 10 ms for E and I, respectively, and membrane capacitance *C* = 500 pF. Neurons emitted spikes when *V(t)* ≥θ, Θ being a relative threshold value set to 20 mV. After the emission of a spike, *V(t)* = *H* for an absolute refractory period of 2 ms and 1 ms for E and I neurons, respectively. All neurons had the same reset potential *H* = 15 mV.

The input current*I* to our LIF neurons was composed of different sources: *I* = *I*_syn_+*I*_ext_+*I*_AHP_. The background current *I*_ext_ modeled the synaptic barrage due to neurons external to the simulated cell assembly. This input is here described as a random train of spikes, distributed in time with Poissonian statistics at a frequency of 2040 Hz and 2400 Hz for E and I neurons, respectively. Each external spike instantaneously depolarizedV(t) by *J*_ext_, a synaptic efficacy randomly chosen from a Gaussian distribution with mean 0.43 mV and 0.56 mV for E and I neurons, respectively, and standard deviation set to the 25% of the mean. To model spike-frequency adaptation [38, 44], excitatory neurons also received an activity-dependent hyperpolarizing current *I*_AHP_ = –*g_c_c*, proportional to a fatigue level *c(t)*, modeling the Ca^2+^ and Na^+^ extracellular concentrations, and with dynamics d*c*/d*t* = –*c/τ_c_* + Σ_k_ δ(*t–t*_k_). Specifically, for each spike emittedat time *t_k_* by the neuron, a unit increase of *c(t)* occurred followed by a decay with constant *τ_c_* = 1 s. This time constant, together withthe unitary currentg*_c_* = 10 pA, was chosen in order to have statistics for Up and Down state durations quantitatively similar to those measured in our experiments.

Current *I*_syn_ was the synaptic input due to the local reverberation of spikes in the modeled cell assembly. Neurons were sparsely connected in the network, having a probability of 20% to be synapticallycoupledto any other neuron. Spikes emitted by neurons in subpopulation B induced in post-synaptic neurons of target pool A an instantaneous polarization of *V(t)* with size randomly extracted once for all from a Gaussian distribution with mean *J_AB_* and standard deviation equal to 25% of the mean. We set *J*_EE_ = 0.43 mV, *J*_EI_ = *J*_II_ = –1.5 mV and *J*_IE_ = 0.56 mV. In order to have slow oscillations only in a subset of 1000 E neurons, the synaptic strength 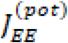 between these neurons was increased to 0.64 mV. To compensate the resulting increase of excitatory current for this subset of neurons, we depressed to 0.36 mV the average synaptic 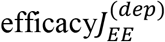 between the subgroups of E neurons active or inactive in the Up state. Emitted spikes were delivered to post-synaptic neurons with a random transmission delay δ set at the beginning of the simulations, and extracted from an exponential distribution with mean values of 22.5 ms and 5.6 ms for E and I presynaptic neurons, respectively. Different timescales aimed to model different timescales for glutamatergic and GABAergic synaptic transmission.

Under mean-field approximation (many neurons sparsely connected and small synaptic efficacies compared to θ–*H*), the network dynamics can be effectively described as a 2D nonlinear system with τ_ν_dν/d*t* = Φ(*c, ν*) – ν and d*c*/d*t* = –*c/τ_c_* + ν [39]. As a good approximation of Φ(*c, ν*), we used the LIF neuron gain function, giving the output firing rate at constant fatigue level *c* and input firing rate ν [29]. The network relaxation timescale τ_ν_ was an arbitrary small time constant, not affecting the nullcline computation (see below). In this theoretical framework, the state variables *c(t)* and *ν(t)* were the fatigue level averaged across neurons and the instantaneous firing rate per neuron for the excitatory pool with Up and Down slow oscillations. In order to consider only this subpopulation of excitatory neurons, we relied on an adiabatic approximation for which Φ(*c, ν*) was computed assuming that the other excitatory and inhibitory pools instantaneously reached their stationary fixed points (Mascaro and Amit, 1999; Mattia et al., 2013). To improve the agreement between such theory and simulations, in computing Φ(*c, ν*) we arbitrarily set the absolute refractory period τ_0_for excitatory neurons to 8 ms, a change thataffected only high-ν activity. For this 2D mean-field system we worked out the nullclines dν/d*t* = 0 and d*c*/d*t* = 0, further distinguishing in the former the stable fixed points with dΦ/dν< 1 from the other points (solid and dashed branches, respectively, in Fig. 5F). Fixed points for the full system were the intersection between the two nullclines, which for the investigated example were stable attractors only when the intersection involved a stable branch of dν/d*t* = 0 (Strogatz, 2001).

#### Modeling DC stimulation

DC fields can be effectively modeled as linearly correlated with the emission threshold θ of the excitatory/pyramidal neurons [45, 46]. This was equivalent to providing an additional ionic current Δ*I* (Fig. 5E), which shifted the asymptotic membrane potential by τ Δ*I/C*, and thus shifting θ up or down. So if λ = Δθ/ΔE was the change of the emission threshold per unit field magnitude, Δ*I*/ΔE = λ C/τ was the modulation factor for the additional ionic current. Here *λ*= 0.12 mm, in the range of values measured in other studies [45, 46].

## Results

Our objective was to determine whether slow and also fast oscillations generated by the cortical network could be up/downregulated by DC electric fields perpendicular to the cortical layers and in that case, to investigate the underlying network mechanisms. This investigation was carried out both *in vitro* and *in silico*.

Unfiltered local field potential (LFP) recordings were obtained from deep layers of a total of 28 ferret neocortical slices (18 visual, 10 prefrontal). All the slices displayed spontaneous slow oscillatory activity consisting of active periods, or Up states, and silent periods, or Down states (Fig. 1C-F, control condition). This is emergent activity from the cortical network that does not require the presence of any pharmacological agent or electrical stimulation, as long as the slices are bathed in ACSF with ionic composition similar to the one *in situ* [23]. The average frequency of the slow oscillations under control conditions was 0.38 ± 0.13 Hz, with an average duration of Up states of 0.53 ± 0.26 s and of Down states of 2.56 ± 1.31 s. During Up states, the local neuronal activity was temporally structured in fast oscillations including those in the beta (15-30Hz) and gamma (30-90Hz) range. Out of the 28 slices, 21 were included in the first part of the study for modulation of oscillations by DC electric fields and 7 were included in the last section where kainate was applied to the bath.

### Electric fields modulate the slow oscillation frequency

To study the effects induced by DC electric fields on the oscillatory activity, we first applied fields of different intensities ranging between -6 and +6 V/m (Fig. 1A). Each intensity (in steps of 1 V/m) was applied for 150 s for each polarity, each one interspersed with 150 s without any DC. The impact of electric fields on the slow oscillations was quantified by estimating the following parameters: SO frequency, duration of Up and Down states, local firing rateduring Up and Down states, coefficient of variation (CV) of the different oscillatory periods and slopes of the transitions from one state to the next one [46].

The control SO frequency was almost doubled with 5 V/m and practically silenced with -5 V/m (Fig. 1C,D). This increase in oscillatory frequency was probably not due to the duration of the Up states during this period since it was hardly modulated by DC fields within the explored intensity range (Fig. 1E,F). In spite of this relative stability of the Up state duration, the same did not occur for the Down state duration.

The increase in SO frequency by increasing DC fields (Fig. 1C) seems to have been a consequence of a reduction in the Down state duration, which was consistent at the population level (Fig. 2). Within the studied range of DC field intensities (−6 V/m to +6 V/m), the positive relationship between oscillatory frequency and DC field intensity was linear in a logarithmically scaled Y axis, hence revealing an exponential relationship (Fig. 2A). Similarly, the parameter of the slow oscillation that was more tightly controlled by the DC field’s intensity was the duration of the Down states, which decreased with increasing DC field intensity (Fig. 2B).

**Figure 2.**
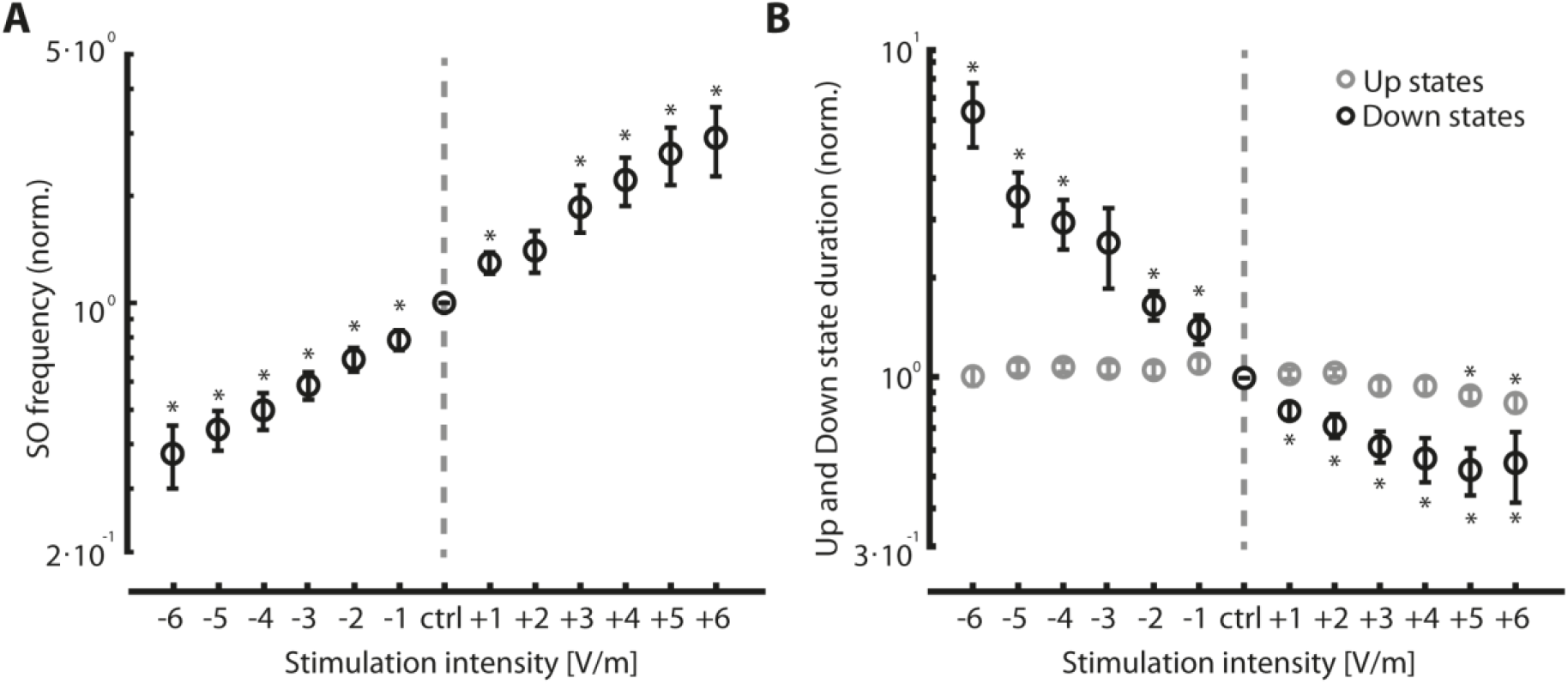
Modulation of slow oscillation frequency and Up/Down state durations by DC fields. A. SO frequency across DC field intensities. One-sample Student *t*-test (*p < 0.05) compared to control condition; n=15 out of 21 slices included in the analysis: -6 V/m, *p*=2·10^-3^; -5 V/m, *p*=3·10^-4^; -4 V/m, *p*=6·10^-6^; -3 V/m, *p*=1·10^-5^; -2 V/m, *p*=1·10^-4^; -1 V/m, *p*=1·10^-3^; 1 V/m, *p*=7·10^-3^; 3 V/m, *p*=1·10^-2^; 4 V/m, *p*=4·10^-3^; 5 V/m, *p*=8·10^-3^; 6 V/m, *p*=2·10^-2^. **B.** Duration of Up states (black) and Down states (gray) across DC field intensities. Same sample size as in A; for Down states: -6 V/m, *p*=4·10^-2^; -5 V/m, *p*=3·10^-2^; -4 V/m, *p*=5·10^-3^; -3 V/m, *p*=7·10^-2^; -2 V/m, *p*=3·10^-3^; -1 V/m, *p*=1·10^-2^; 1 V/m, *p*=8·10^-4^; 2 V/m, *p*=4·10^-4^; 3 V/m, *p*=8·10^-5^; 4 V/m, *p*=3·10^-4^; 5 V/m, *p*=2·10^-4^; 6 V/m, *p*=1·10^-2^; for Up states: 5 V/m, *p*=2·10^-2^; 6 V/m, *p*=2·10^-2^. All data were normalized to the control condition, averaged across slices, and are shown as mean ± S.E.

More specifically, the decrease in Down state duration by DC fields of increasing intensity was linear in most neurons when represented in a logarithmic scale (see Fig. 3A for a single example). The slope of this relationship showed heterogeneity across slices when the whole population was considered (Fig. 3B). In order to quantify this heterogeneity, we defined a Modulation Index as the slope of the linear fit to the relationship between the logarithm of Down state duration and the DC field intensity (i.e. the slope for each regression line in Fig. 3B) (Fig. 3B, inset). In 6 out of 21 slices, the Modulation Index was lower than 0.05 (Fig. 3B, inset). We excluded those slices from the population level analysis (Fig. 2) considering that DC field did not modulate slow oscillatory frequency in those slices (Fig. 3B, gray lines). We further explored why some slices were hardly modulated by an electric field and we found that it depended on the initial oscillatory activity of each slice in control conditions: the Modulation Index correlated significantly with the absolute value of the Down state duration during control conditions (Fig. 3C; multiple linear regression, p=4·10^-6^) such that the longer the Down state duration in control conditions, the larger the Modulation Index. Furthermore, we detected a weak, yet significant, correlation between the Modulation Index and the duration of the Up state during control conditions (Fig. 3C, inset; multiple linear regression, p=0.01). These results indicate that slices with an intrinsic high level of neuronal excitability (i.e. short Down state duration and high SO frequency: *circa* 0.5 Hz) were less susceptible to the further increase in excitability as induced by the DC fields and therefore no modulation of the SO frequency was observed.

**Figure 3.**
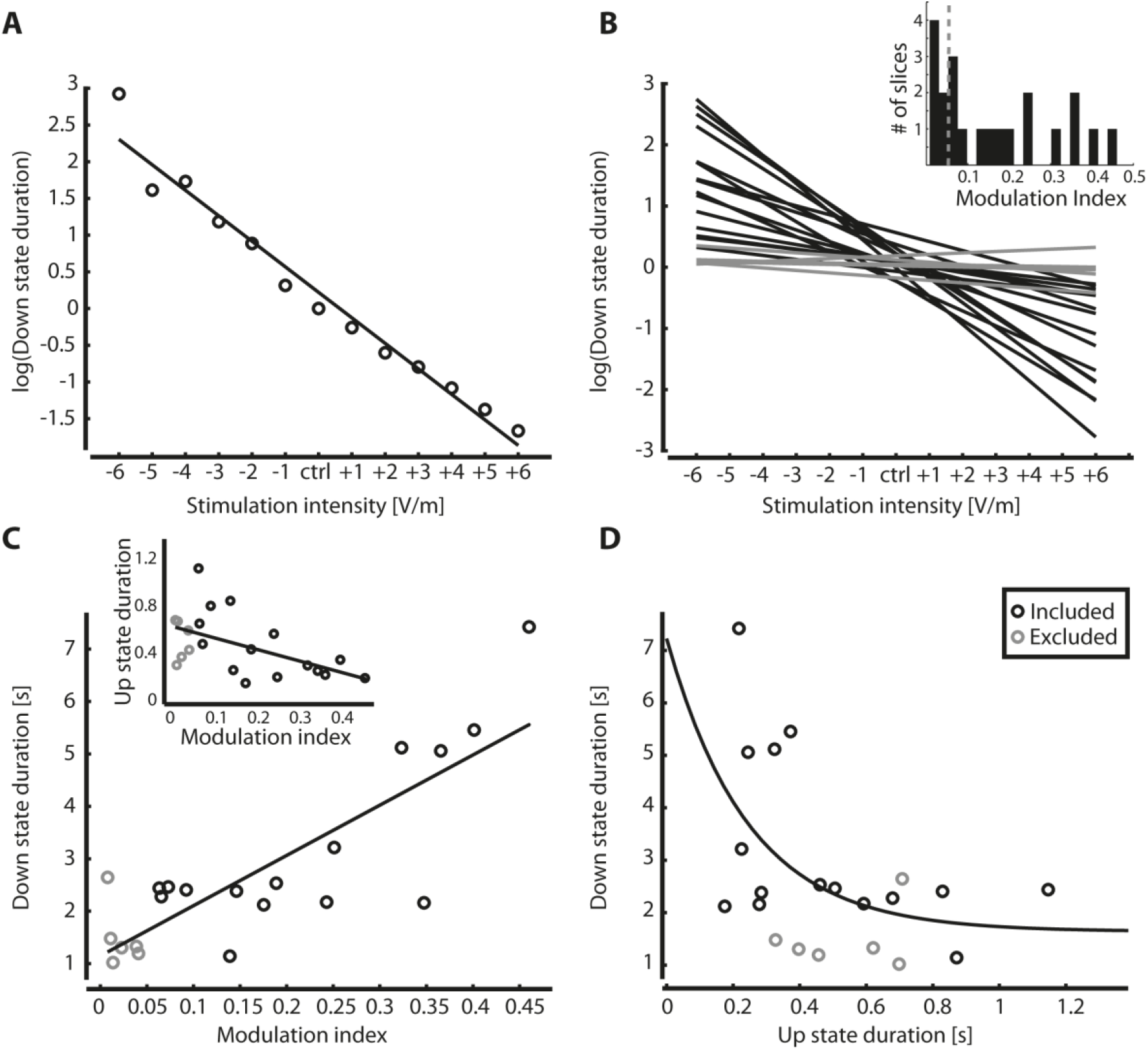
Relationship between duration of Down states and the modulation of slow oscillations by DC fields. **A.** Linear correlation between the logarithm of the values of Down state duration and the DC field intensity (representative case). Multiple linear regression, *p*=2· 10^-9^. **B.** A Modulation Index for the effects of the DC field on the slow oscillation was obtained for each slice as the slope of the linear fit obtained as shown for a representative case in A. Recordings with a Modulation Index lower than 0.05 (gray lines) were considered non-modulated and were therefore excluded from the population analysis (n=6 out of 21 excluded, see inset). **C**, Linear correlation between the Modulation Index and the Down state duration (and with the Up state duration in the index) for all the 21 recordings. Multiple linear regression, *p*=4·10^-6^ for Down state duration, *p*=1·10^-2^ for Up state duration. Gray circles represent the 6 excluded cases with Modulation Index < 0.05. **D.** Exponential correlation between Up states duration and Down states duration. Nonlinear exponential regression, MSE=2.37.

Furthermore, we studied how the relationship between Up and Down state durations influenced the modulatory effect of electric fields and found a negative exponential relationship between the duration of Up and Down states (Fig. 3D; nonlinear exponential regression, mean squared error= 2.37), as theoretically predicted [29].

### Up states remain stable in the presence of electric fields

As mentioned above, Up states appeared to be very robust in the presence of DC fields regarding duration (Fig. 1C) and firing rate (Fig. 1F). Upon closer inspection, Up state duration remained stable with increasing DC field although there was a small but significant decrease at the maximum positive intensities (+5 and +6 V/m; Fig. 2B), and the relative firing rate remained mostly constant (Fig. 4A). Maintaining stable Up states while modulating Down states suggests that there could be a remarkable dynamic uncoupling between Up and Down states. In other cases of perturbation of the cortical network such as by varying inhibition levels [46] or changing temperature [42], the resulting modulation of both Up and Down states’ duration was not independent but dynamically related. For example, decreasing inhibition caused progressively shorter Up states of increasing firing rate followed by increasingly longer Down states [47]. In the range of DC fields that we have been working with here, the modulation is centered in the Down states and does not affect Up states, but we cannot rule out that DC fields of larger amplitudes would exert a modulation of Up state’s duration and amplitude.

**Figure 4.**
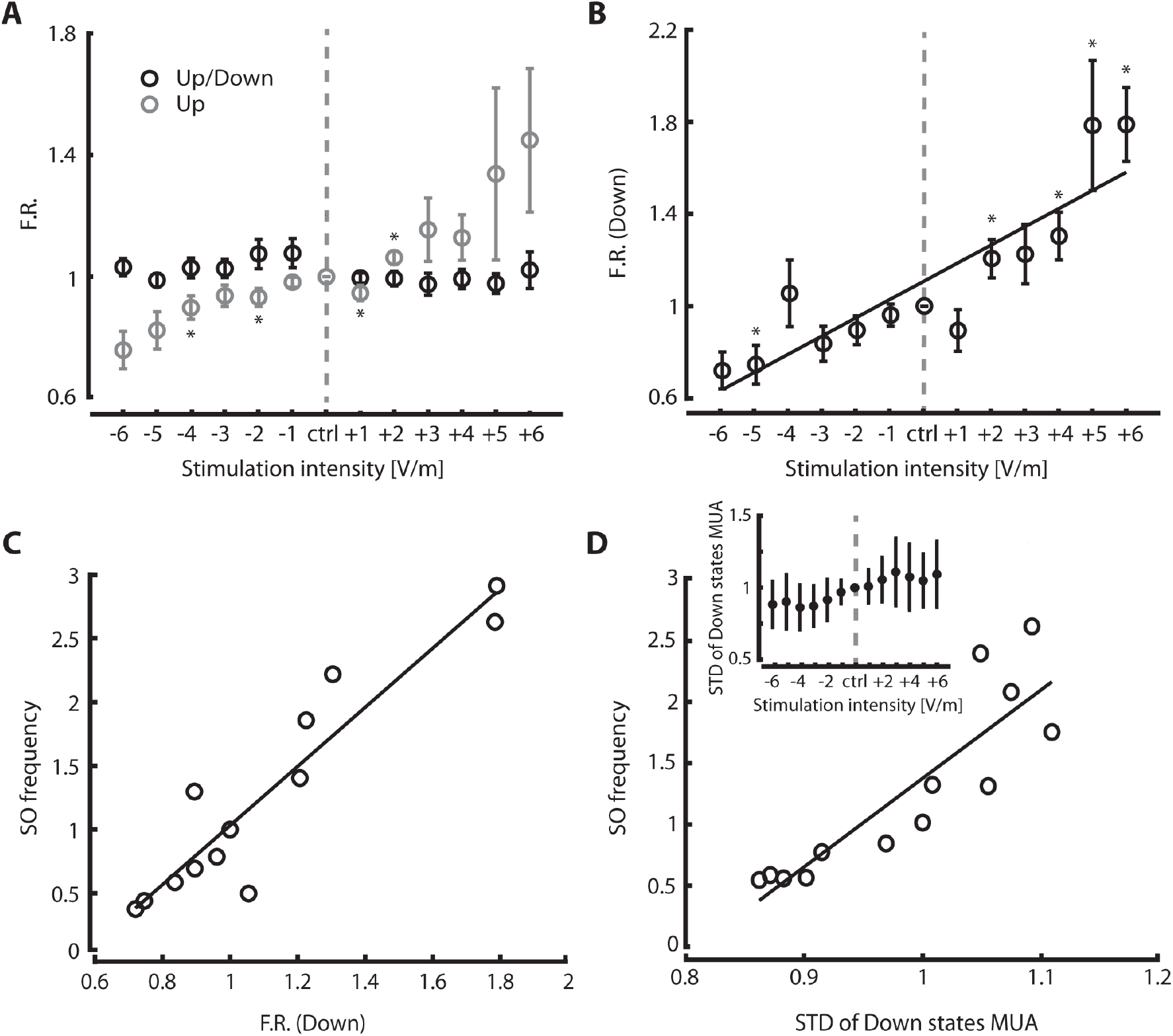
Modulation of firing rate in Up and Down states by DC fields. **A.** Relative firing rate (black) and absolute firing rate (gray) during Up states across stimulation intensities respect to control condition (* p<0.05); n=15 out of 21 slices included in the analysis; one-sample Student *t*-test: -4 V/m, *p*=4 10^-2^; -2 V/m, *p*=4·10^-2^; 1 V/m, *p*=4·10^-2^; V/m, *p*=8·10^-3^. **B.** Firing rate during Down states across stimulation intensities. Same sample size as in A: -5 V/m, *p*=4.7·10^-2^; 2 V/m, *p*=3·10^-2^; 4 V/m, *p*=1·10^-2^; 5 V/m, *p*=2·10^-2^; 6 V/m, *p*=1·10^-3^). **C.** Linear correlation between the absolute firing rate during Down states and the SO frequency. Multiple linear regression, *p*=3·10^-6^. **D.** Linear correlation between the standard deviation of log(MUA) during the period between 250 ms and 50 ms before the following Up state was triggered (shown in the inset across DC field intensities), and the SO frequency. Multiple linear regression, *p*=8·10^-5^. All data were normalized to the control condition, averaged across slices, and are shown as mean ± S.E.

### Electric fields modulate firing rate during Down states

Given this dynamic uncoupling between Up and Down state modulation by DC fields, the following question naturally arises: what are the mechanisms by which DC fields modulate Down state durations and consequently oscillatory frequency? Downstates are relatively silent and DC fields modulate network excitability and the increase in excitability translates into an increase in firing rate also during Down states, therefore “silent” states become less silent. This resulted in a situation where the relative firing rate (Up *versus* Down state firing rate) remained rather constant for the whole range of electric fields (from -6 to +6 V/m; Fig. 4A). However, while the relative firing rate was stable, the absolute firing rate during both Up (Fig. 4A) and Down states (Fig. 4B) decreased for negative fields and increased for positive fields (see figure legend for statistics). Even when Down states are defined as silent periods in between Up states, in the cortical slices there is a certain amount of firing, in particular in infragranular or deep layers [23], which is associated with the initiation of Up states (see Fig. 1 in Compte *et al*., 2003). The origin of Up states in deep layers is also a consistent finding *in vivo* [41, 48]. More specifically, neuronal firing rate in deep cortical layers during Down states increased with positive DC fields (Fig. 4B), which in turn was linearly correlated with the SO frequency (Fig. 4C). This may suggest that the neuronal firing during Down states linearly controls the initiation of the subsequent Up state, which is also theoretically supported by our computer model of the network (see below). We could also estimate the network firing rate as the standard deviation of the relative firing rate during Down states, which was also modulated by the DC fields and correlated with the SO frequency (Fig. 4D). These observations provided a mechanistic hint into how the exponential regulation of Down state duration is achieved, and that is by increasing the firing rate and its fluctuations, and thus the probability to start a new Up state.

Once the Up state has been triggered, its duration and firing are resilient to depolarizations/hyperpolarizations induced by DC electric fields. This observation further supports the hypothesis that depolarization induced by DC fields is negligible in size with respect to the changes of the membrane potential due to incoming synaptic activity during Up states. It would also explain why the only remarkable changes can just be measured during Down states.

It seems reasonable to think that larger DC fields of any sign could have an effect on the Up state duration and firing rate, given that each V/m just corresponds to a neuronal membrane depolarization of 0.5 mV [18]. However, due to the methodological constraints mentioned above, we were not able to explore the effect of DC fields of larger amplitudes.

### DC field modulation of slow oscillation dynamics in a cell assembly model

So far, we have proved that slow oscillations in cortical slices are strongly affected by the application of uniform and temporally constant DC fields. DC fields are known to modulate the excitability level of single pyramidal neurons by polarizing the membrane potential [18, 45, 46]. On the other hand, slow oscillations are a collective phenomenon mainly driven by the synaptic interaction between a large number of neurons [23, 40, 49]. Thus, the modulation of slow oscillations by DC fields should result from the interplay of these two microscopic and macroscopic scales. To further investigate the mechanisms underlying such multi-scale phenomenon we devised a cell assembly model displaying slow oscillations. Modeling MUA from simulated spiking activity (Fig. 5A) allowed us to perform the same analysis used for *in vitro* recordings such as detecting transitions between Up and Down states. Network parameters were chosen to reproduce rasters of logarithmically scaled MUA (Fig. 5B) and state duration statistics (Fig. 5C) quantitatively similar to what was experimentally measured (Fig. 1). Slow oscillations emerged from the oscillatory compensation between two opposite nonlinear forces: a self-excitatory net feedback due to the synaptic reverberation of emitted spikes, which rapidly drove the system activity to the high firing rates of the Up states, and an activity-dependent fatigue level determining self-inhibitory K^+^ currents underlying Up state termination, related to Ca^2+^ and Na^+^ intracellular concentrations [38, 44]. With mean field approximations, the network dynamics can be effectively represented in a two-dimensional space as a relaxation oscillator [29], with state variables ν(t) and c(t) associated respectively to the firing rate of the excitatory pool with Up/Down oscillations and the average fatigue level among the same neurons (Fig. 5A top). In this (c, ν) plane, the network state moved in an orbit (Fig. 5D) in which Up and Down states were the relaxation phases following the stable branches of the nullclinedν/dt = 0. Up (Down) states terminated when the right (left) knee of this nullcline was reached and a fast downward (upward) transition occurred (see Materials and Methods for more details).

**Figure 5.**
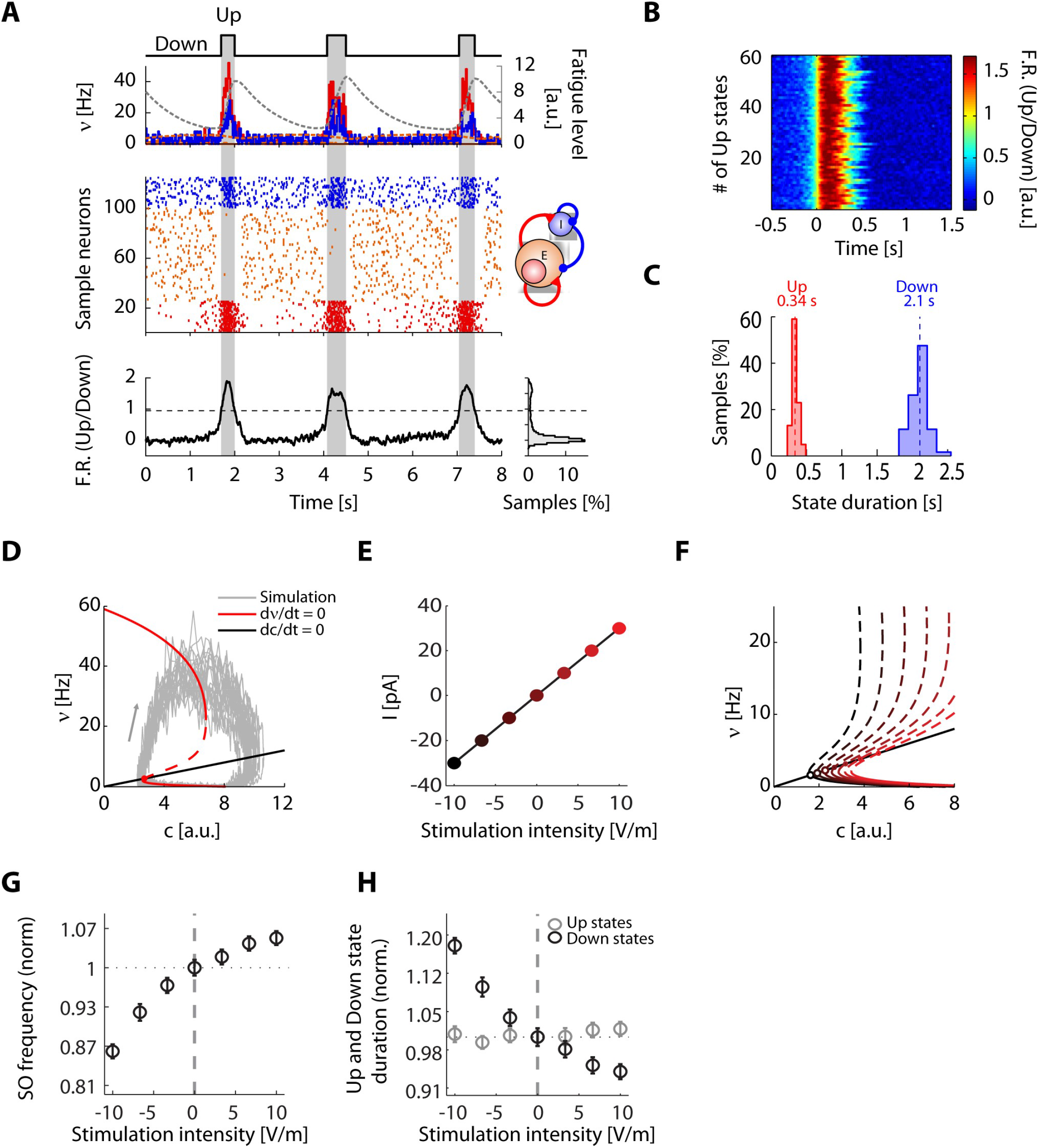
Structure, simulation and predictions of a cell assembly model. A. A simulated network composed of 80% excitatory nodes (neurons) and 20% inhibitory nodes generated slow oscillations, characterized by the typical modulation of firing rate (top row, red (excitatory) and blue (inhibitory) lines). Ending of the Up states in the model was controlled by a mechanism of fatigue (top row, gray dotted line) that then decreased during the Down state. Relative firing rate (bottom row) was obtained as the sum of the instantaneous firing (central row). **B.** Relative firing rate after alignment of all the Up states occurring in 100s of simulation. **C.** Typical duration of Up and Down states in the model. **D.** In the (c,ν) space, the network moved in an orbit (gray lines represent orbits obtained in the simulation) in which Up and Down states were the relaxation phases following the stable branches of the nulldmedv/dt=0 (red line, stable where the line is continuous). **E.** Amount of variation in the baseline ionic current used to simulate DC fields. **F.** The simulation of DC fields provoked a shift of the nullclines in the (c, ν) space. **G.** Predicted change in the SO frequency. **H.** Predicted change in the Up (gray circles) and Down (black circles) state durations.

To implement in the model the modulation of the neuronal excitability by DC fields, we changed the baseline ionic current for the excitatory neurons by an amount *ΔI* proportional to the DC field intensity (Fig. 5E). Changing *ΔI* in response to a DC field application induced a horizontal shift of the ν-nullcline (Fig. 5F). As a result, the intersection between c- and ν-nullclines (open circles, fixed-points of the 2D-system) moved towards lower firing rates as the negative field intensity increased, thus making the network less excitable and the Down state a preferred state, i.e. longer Down states and shorter Up states [29], as in our experimental findings (Fig. 3D). In our simulations, this decrease inexcitability induced a monotonic decrease of the SO frequency (Fig. 5G), which was mainly due to the elongation of Down state durations (Fig. 5H). All these trends were remarkably similar to those measured and shown in Fig. 2. As for the experiments, Up state durations were almost unaffected by DC field changes.

### Negative DC field intensities decrease high frequency power after kainate application

Back to our *in vitro* experiments, we next quantified the potential modulation by DC field so high frequencies in the beta (15-30 Hz) and gamma (low 30-60, and high 60-90 Hz) ranges that are generated by the cortical circuit during the Up states [20, 50, 51]. This quantification was carried out in 17 out of the 21 slices included in the previous sections, since in the remaining 4 slices the high frequency component was not stable over time.

In most slices (13 out of 17) there was no modulation of beta and gamma frequencies by neither positive nor negative DC fields, although some did show modulation (Fig. 6A,B). We reached this conclusion by defining a Modulation Index that was calculated separately for beta and gamma frequencies (Fig. 6C). The Modulation Index was calculated as the slope of the linear fit to the relationship between DC field intensity (V/m) and beta/gamma band power (see Methods). Similar to the analysis on the slow oscillations, we considered that a Modulation Index under 0.05 corresponded to a lack of modulation for both the beta and gamma ranges. By doing this, we detected that DC fields modulated emerging beta and gamma power in 5 and 4 out of the 17 slices, respectively. In these slices there was both a decrease in power by negative DC fields and an enhancement by positive DC fields (Fig. 6C).

**Figure 6.**
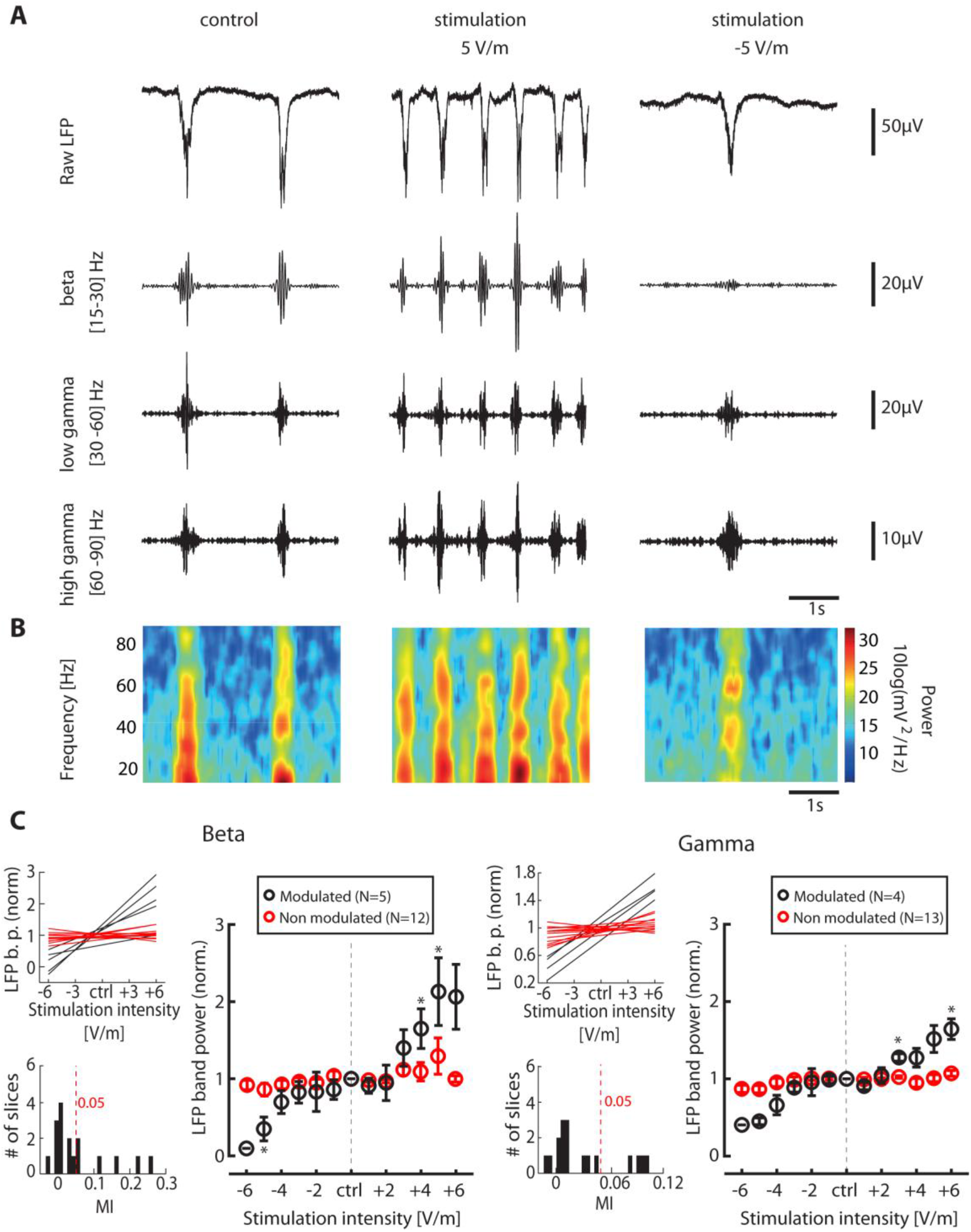
Modulation of activity in the beta and gamma frequency ranges. A. Analysis of changes in high frequencies by the study of LFP raw traces of a sample recording. Recordings (4 s) are shown for 3 different conditions: control (left), positive DC field with intensity of 5 V/m (center), negative DC field with intensity of 5 V/m (right). Traces were band-pass filtered in different ranges: beta (15-30 Hz), low gamma (30-60 Hz), high gamma (60-90 Hz). **B.** Spectrogram of LFP in each condition for the same time periods illustrated in panel A. **C.** Modulation of beta (left) and gamma (right) activity across DC field intensities at the population level, measured respectively as the averaged power in the frequency ranges 15-30 Hz and 30-90 Hz. Individually for beta and gamma, the slices were divided into two groups based on a Modulation Index (bottom insets). Recordings with a Modulation Index <0.05 were considered non-modulated by the stimulation and were therefore excluded from the analysis. One sample Student t-tests (*p < 0.05) comparing to control condition: n=5 and n=4 out of 17 slices modulated at beta and gamma, respectively: beta: -5 V/m, *p*=4·10^-2^; 4 V/m, *p*=3·10^-2^; 5 V/m, *p*=4·10^-2^; gamma: at 3 V/m, *p*=5·10^-3^; 6 V/m, *p*=2·10^-2^. MI, Modulation Index.

The local network in cortical slices can generate beta and gamma frequencies during Up states in the absence of any additional neurotransmitter or electrical stimulation [51], although the power in the beta/gamma range is lower than in the cortex *in situ* [50, 52]. Kainate is a glutamate receptor agonist that has been often used in the hippocampus [53] and cortex *in vitro* [54] in order to enhance high frequencies. Replicating previous studies in the entorhinal cortex [54], we found that kainate application in the neocortex terminated spontaneous Up and Down state dynamics and shifted the network into a state of continuous firing (Fig. 7A). More specifically, kainate increased the beta and gamma power in 7 out of 7 cases (Fig. 7). For instance, a representative slice showed an overall increase in the high frequency range cause by kainate, peaking at 32 Hz in this case (Fig. 7B), and peaking at 30 Hz in the population (data not shown).

**Figure 7.**
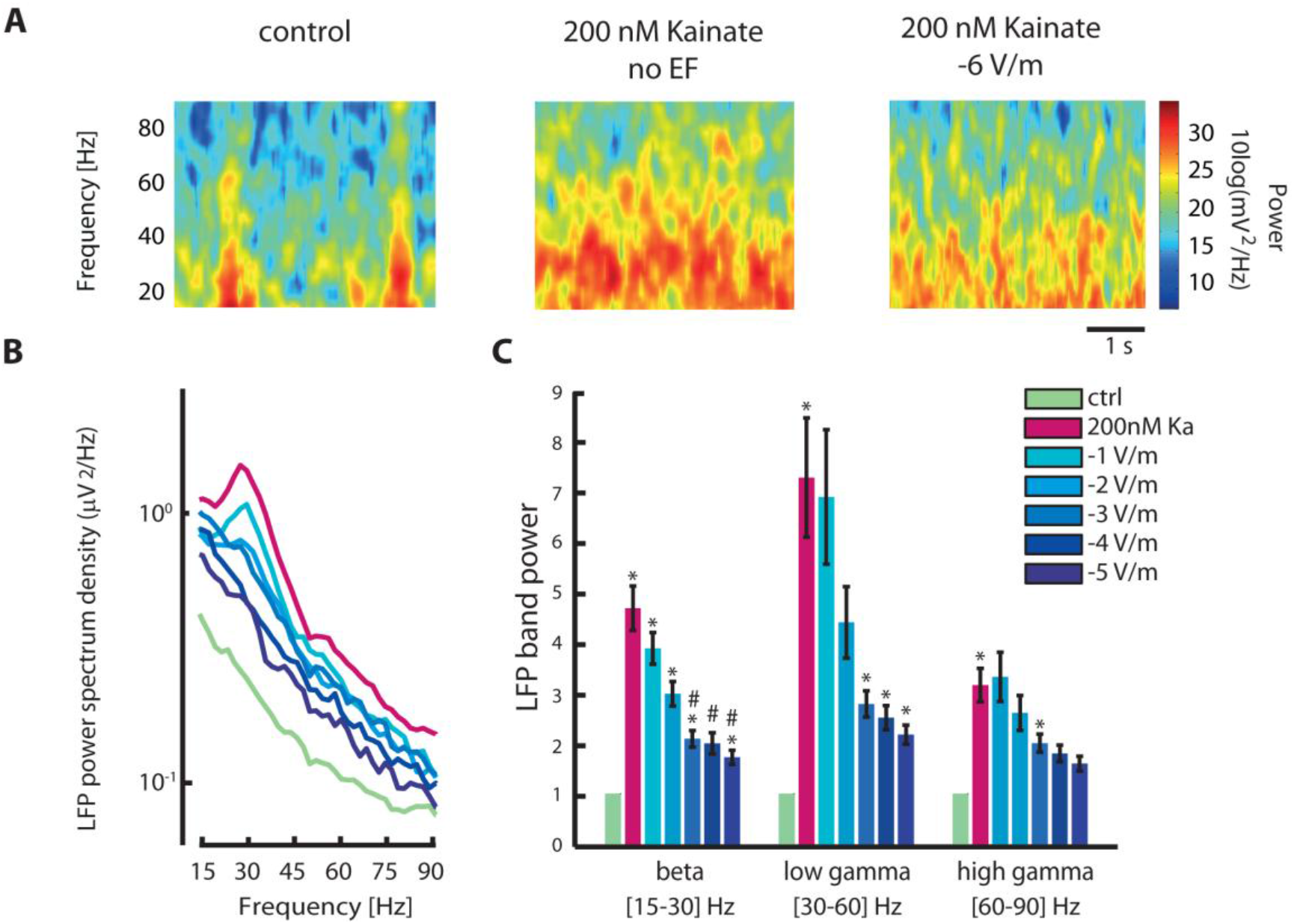
Effects of DC fields on high frequencies after increasing activity with kainate. **A.** Spectrograms of 4 s of a representative recording in 3 different conditions: control (left), control after bath application of 0.2 μM kainate (Ka) (center), negative DC field of 5 V/m (right). **B.** LFP power spectrum density in all conditions (color coded) of the same sample recording. **C.** Mean power changes in beta (15-30 Hz), low gamma (30-60 Hz) and high gamma (60-90 Hz), for all conditions. Data were normalized to the control condition and then averaged across slices. One-sample Student *t*-test (*p < 0.05) compared to control condition; n=7 slices: beta: afterKa, *p*=9·10^-3^; -1 V/m, *p*=6·10^-3^; -2 V/m, *p*=1·10^-2^; -3 V/m, *p*=2·10^-2^; -5 V/m, *p*=4·10^-2^; low gamma: afterKa, *p*=4.5·10^-2^; -3 V/m, *p*=2·10^-2^; -4 V/m, *p*=3·10^-2^; -5 V/m, *p*=3·10^-2^; high gamma: afterKa, *p*=2·10^-2^; -3 V/m, *p*=4•10^-2^. *#p* < 0.05 compared to kainate condition; two-sample Student*t*-test: beta: -3 V/m, *p*=3·10^-2^; -4 V/m, *p*=3·10^-2^;-5 V/m, *p*=3·10^-2^.

Positive DC fields (0 to +6 V/m) did not induce any further increase in high frequencies once kainate had been applied, suggesting that the isolated cortical network under the influence of kainate reached a saturation of its capability to generate high frequency activity while in the absence of other neurotransmitters. Negative DC fields, on the other hand, removed the emergent peak at 32 Hz induced by kainate and reduced the overall activity in the beta and gamma ranges approaching the control level (Fig. 7B). Averaged results across slices showed that, after the application of kainate, the power was largely increased with respect to the control condition, with a stronger effect on the beta and low gamma ranges (Fig. 7C). Negative DC fields after kainate application significantly reduced the power with respect to the kainate condition in all studied frequency bands (Fig. 7 C).

## Discussion

We have studied the effects of electrical DC fields of varying intensities and direction (inward/outward) on the emergent neural activity of ferret cortical slices. We found that augmenting DC positive field intensity increased the frequency of slow oscillations by decreasing the duration of Down states while Up state durations remained relatively stable, although they shortened for the greatest field intensities (+5-6 V/m). The shortening of Down states showed a positive correlation with the firing rates in Down states, thus enhancing the probability to initiate the subsequent Up state. This mechanistic explanation suggested by the experimental observations was quantitatively proved in a computer model. We reproduced the experimental findings in a cell assembly model whose collective dynamics are fully captured by mean-field description and where the stability of Down states is the prevailing mechanism underlying the modulation of emergent activity. Augmenting DC positive field intensity increased in few cases the power of beta/gamma frequency bands (occurring during the Up states of the slow oscillations), whereas augmenting DC negative field intensity was highly effective to parametrically decrease the power of beta/gamma frequency bands, being able to practically cancel the effect of kainate on high frequencies. Our findings have implications for the understanding of cortical oscillations and the mechanisms by which they are modulated by DC fields, and they may contribute to the future development of more specialized tools to treat neuropsychiatric conditions.

We addressed the effect of long-duration (150 s) electric fields stimulation on the neocortical activity by applying weak (−6 to +6 V/m) uniform electric fields to ferret active cortical slices *in vitro* that exhibited spontaneous slow rhythmic activity [23]. This activity is highly similar to slow wave sleep and deep anesthesia [55] and has been proposed to be a default emergent activity of the cortical network [22]. The *in vitro* preparation provided a manageable cortical network, facilitating the understanding of the mechanisms underlying the exogenous modulation of the cortical network. The electric field strength was limited by the duration of the stimulation sessions (150 s): such long-lasting pulses, and depending on the intensity of the currents, could affect the Ag/AgCl electrodes by varying their mass–alternating polarities were used to prevent this-. Other authors using higher intensities have stimulated for shorter periods (0.5 s) [12]. To prevent variations in the intensity of the field and high frequency noise due to deterioration of the electrodes, we were limited to fields between a maximum range of - 6 to +6 V/m. In any case, it has been demonstrated in previous studies that 1 V/m is about the minimum intensity needed to modulate network activity [27] and 2-4 V/m exogenous fields are enough to modulate slow oscillations *in* vitro[18]. Furthermore, the range of electric fields estimated to act on the cortex during tDCS in humans is often <1 V/m [24, 56].

### Mechanisms of modulation of slow oscillatory frequency

When an electric current is directly applied to the cortex, the area underneath the anode is depolarized, while the neurons below the cathode are hyperpolarized [57]. Previous studies in hippocampus slices [12, 58] demonstrated that constant positive DC fields cause a somatic depolarization of about 1 mV for each applied V/m in pyramidal neurons, while negative fields induce an equivalent hyperpolarization of the membrane in the same neural population [13, 32]. Such changes result from local differences in the extracellular and intracellular distribution of ionic charges in response to the imposed extracellular voltage gradient. These small changes in the neuronal membrane potential have been shown to have a more noticeable impact when the activity resulting from the whole network is considered, both in cortex [18] and hippocampus [15]. Looking into slow oscillations, we observed a shortening of the Down states with the application of positive fields, which followed the depolarization of the membranes, therefore bringing the cells nearer to the threshold of Up state production. This increase in the excitability of single neurons would contribute to the increase in fluctuations of the network firing rate (Fig. 4D), resulting in a higher probability of producing Up states. On the other hand, the membrane of excitatory neurons is hyperpolarized under the application of negative fields, hence lowering the amplitude and fluctuations of the population firing rate and in turn reducing the probability of Up state production, resulting in an elongation of the Down state duration in our experiments. In other words, the changes in single-neuron excitability affect the collective dynamics of the probed cell assembly turning out to be an effective modulation of the Down state stability. In the model, this stability modulation which made the Down state a more preferred or a less preferred attractor state allowed to fully reproduce the features of the *in vitro* slow oscillations we measured.

Remarkably, the average duration of the Up states remained relatively constant across stimulation intensities. This effect could be associated with the small change in membrane polarization induced by DC fields in the range of the intensities applied, which could be nonetheless enough to modulate the activity during Down states owing to the amplification carried by the network. Up states are defined as *high conductance* states [59–62]: during the Up states there is an increase in the membrane conductance that leads to a three- to five-fold decrease in input resistance. This would further reduce the effect of the DC fields on the membrane which, within the intensity range that we tested, could modulate the Up state spontaneous firing, but not their duration.

But to what extent can Down states vary independently of Up states? In a study where progressive blockade of inhibition causes a correlated increase in firing rate during Up states, shortening of Up states occurred concurrently with elongation of Down states [47]. Therefore, Up and Down state durations were dynamically linked, since the firing rates of Up states determined the duration of the subsequent Down states through the activation of neuronal hyperpolarization. However, in the current study we observed that Up and Down duration variations can be uncoupled and modulate Down state duration while leaving Up states almost unaffected. We can discuss several possible causes at play. First of all, as mentioned above, DC fields increases or decreases network excitability, thus affecting the firing during both Up and Down states. Therefore, the “relative” firing rate (Up/Down) remains remarkably constant (Fig. 4A) while it parametrically increased with the inhibition blockade. In the same direction, modulating synaptic inhibition is restricted to periods with presynaptic firing (Up states) while it has low effect in Down states (i.e. low or no firing).

It is interesting to notice how this mechanism of dynamic uncoupling between Up and Down states is actually dependent on the initial network dynamics. In fact, when the initial oscillatory state is characterized by longer Up states and shorter Down states, then the Up/Down dynamics coupling is more difficult to break and the electrical stimulation becomes less effective (Fig. 3) because a network in this area of operability is a network with higher excitability that spends more time on active discharging states, and that has in turn smaller room for modulation.

### Mechanisms of modulation of high frequency activity

Both the small amplitude of hyper/depolarization induced by DC fields, the excitatory / inhibitory balance and the high conductance state can be at the base of the only slight modulation of the activity in the beta and gamma ranges. In most cases the activity at those frequencies remained unchanged across stimulation intensities and orientations. We have not identified factors that can explain why some slices showed no modulation at all while the other ones did. Nevertheless, we ruled out the relationship of high frequency modulation with network firing rate, Up/Down duration and frequency of the slow oscillations. Given that there is a large interest in the potential cognitive impact of weak tDCS [63, 64] and that cognitive performance is associated with gamma frequencies [65–68], we consider that this specific issue requires further research in particular with a broader range of intensities and in the alert brain.

Interestingly, in the cases in which there was high frequency modulation, we observed a stronger increase in the beta activity than in the gamma activity with the application of depolarizing fields. Such frequency-specific modulation could arise from changes in the network excitability by the electric stimulation, for which we suggest the following explanations. The observed increase in the Up state firing rate provoked by positive electric fields would promote synaptic reverberation, i.e. a state in which recurrent connections among excitatory and inhibitory neurons stimulate firing of all types of neurons. Although inhibitory cells are fewer in number and due to their morphology less likely to be directly affected by constant electric fields [46], they are also recruited through the recurrent connections with excitatory neurons. An increased activation of the excitatory cells with positive fields would in turn increase the recruitment of the inhibitory cells and, given the role of the latter in the generation of high frequency oscillations [51], this overall increase in activity would result in the production of beta and gamma rhythms. Negative field stimulation would instead prevent the network from reaching the resonant state, with a consequent decrease in the overall frequency spectrum. This is in fact strikingly demonstrated in the case of pharmacologically enhanced activity (Fig. 7). The application of kainate [69], used as means to confirm that our slices were indeed able to produce high frequency rhythms - given the low proportion of slices whose high frequencies responded to electric fields-, induced a strong depolarization of the network (with emergent peaks across beta and gamma) that resulted in an almost continuous Up state. Negative fields were able to decrease the altered network activity almost back to the initial condition, thus cancelling the effect of kainate.

Finally, it is interesting to remark that the DC-induced modulation of the Down state stability can have an impact on the way in which slow oscillations are shaped by the connectivity structure and the heterogeneity of the underlying cortical network. Indeed, recent findings have highlighted how the variability of these slow rhythms [70] and the reproducibility of some features of the travelling Down-to-Up wavefronts spontaneously expressed by the slices [71] strongly depend on the stability of the Down state of local cell assemblies. More specifically, the laminar structure of the cortex determines the shape of the activation wavefronts only when the Down state is marginally stable, as in the case we described here as control condition (null DC field). If this happens, cortical layers 4 and 5 (those with higher firing rates) lead such wavefront propagation. This is not the case if the slice excitability is smaller or higher compared to the control condition. With respect to this, DC field can then be exploited to guide the excitability level of the network towards an optimal condition for the Down state stability which makes the network maximally sensitive to the underlying structure, and thus compensating perturbations of the tissue excitability expressed in some neurological disorders or lesions.

## Conclusions

This study demonstrates that relatively small electric fields of similar amplitude to the ones measured *in vivo* [24] are enough to impose a complete control of the frequency of slow oscillations generated by the cerebral cortex *in vitro*. This control occurs by regulating the excitability of neurons during the Down states, resulting in a precise control of their duration and thus of the frequency of the oscillations which we quantitatively explore in a computer model. These same electric fields are not powerful enough though to modulate the duration of the Up states, suggesting that the intrinsic network controls are not surpassed by the exogenous fields within the field intensity range that we used. Finally, we demonstrate that decreasing excitability by means of negative electric fields is enough to radically decrease beta and gamma synchronization during Up states.

## Acknowledgments

This work was supported by the Spanish Ministerio de Ciencia e Innovación (BFU2014-52467-R) and by PCIN-2015-162-C02-01 (FLAG ERA) to MVSV and WaveScales(#604102; Human Brain Project) to MVSV and MM. IDIBAPS (MVSV) is funded by the CERCA Programme (Generalitat de Catalunya). We would like to thank Dr. Cristina Gonzalez-Liencres for her comments on the manuscript.

